# Genome-wide CRISPR screen reveals host genes that regulate SARS-CoV-2 infection

**DOI:** 10.1101/2020.06.16.155101

**Authors:** Jin Wei, Mia Madel Alfajaro, Ruth E. Hanna, Peter C. DeWeirdt, Madison S. Strine, William J. Lu-Culligan, Shang-Min Zhang, Vincent R. Graziano, Cameron O. Schmitz, Jennifer S. Chen, Madeleine C. Mankowski, Renata B. Filler, Victor Gasque, Fernando de Miguel, Huacui Chen, Kasopefoluwa Oguntuyo, Laura Abriola, Yulia V. Surovtseva, Robert C. Orchard, Benhur Lee, Brett Lindenbach, Katerina Politi, David van Dijk, Matthew D. Simon, Qin Yan, John G. Doench, Craig B. Wilen

## Abstract

Identification of host genes essential for SARS-CoV-2 infection may reveal novel therapeutic targets and inform our understanding of COVID-19 pathogenesis. Here we performed a genome-wide CRISPR screen with SARS-CoV-2 and identified known SARS-CoV-2 host factors including the receptor ACE2 and protease Cathepsin L. We additionally discovered novel pro-viral genes and pathways including the SWI/SNF chromatin remodeling complex and key components of the TGF-β signaling pathway. Small molecule inhibitors of these pathways prevented SARS-CoV-2-induced cell death. We also revealed that the alarmin HMGB1 is critical for SARS-CoV-2 replication. In contrast, loss of the histone H3.3 chaperone complex sensitized cells to virus-induced death. Together this study reveals potential therapeutic targets for SARS-CoV-2 and highlights host genes that may regulate COVID-19 pathogenesis.

## Introduction

Severe Acute Respiratory Syndrome-Coronavirus-2 (SARS-CoV-2), the causative agent of Coronavirus Disease 2019 (COVID-19), represents the greatest public health threat in a century. More than 7,500,000 people have been infected with more than 420,000 deaths globally (*1*). Novel therapeutics and vaccines are desperately needed. Coronaviruses are enveloped, positive-sense RNA viruses with genomes of approximately 30 kb that exhibit broad host-range among birds and mammals and are typically transmitted via the respiratory route (*2, 3*). There are four circulating seasonal coronaviruses in humans (NL63, OC43, 229E, and HKU1) and three highly pathogenic zoonotic coronaviruses (SARS-CoV, MERS, and SARS-CoV-2), none of which have effective antiviral drugs or vaccines (*4–7*).

Viral entry, the first stage of the SARS-CoV-2 life cycle, is mediated by the viral spike protein. The receptor binding domain of spike binds to the cell surface receptor angiotensin-converting enzyme 2 (ACE2), a major determinant of host range and cell tropism (*8, 9*). The coronavirus spike protein requires two proteolytic processing steps prior to entry. The first cleavage event occurs at the interface of the S1 and S2 domains of the spike protein (*10, 11*). This can occur in the producer cell, the extracellular environment, or in the endosome and can be mediated by several proteases including furin and the plasma membrane protease TMPRSS2 (*12–14*). A second proteolytic event is required within S2 to expose the viral fusion peptide and enable membrane fusion. This second cleavage event can occur at the target cell plasma membrane by TMPRSS2 or in the endosome by Cathepsin L (*14, 15*). Upon viral membrane fusion, the viral RNA is released into the cytoplasm where it is translated and establishes viral replication and transcription complexes before assembling and budding (*16–18*). The host genes that mediate these processes largely remain elusive.

Identification of host factors essential for infection is critical to inform mechanisms of COVID-19 pathogenesis, reveal variation in host susceptibility, and identify novel host-directed therapies, which may have efficacy against current and future pandemic coronaviruses. To reveal host genes required for SARS-CoV-2 infection and cell death, we performed a genome-wide CRISPR screen in a *Chlorocebus sabaeus* (African green monkey or vervet) cell line, Vero-E6. Surprisingly, although SARS-CoV-2 is an RNA virus that replicates in the cytosol, our screen revealed an abundance of host genes that function in the nucleus. Specifically, we identified the SWI/SNF chromatin remodeling complex, key TGF-β signaling components, and the alarmin HMGB1 as pro-viral while we revealed the Histone H3.3 complex as anti-viral. We individually validated 25 of the CRISPR gene hits and demonstrated that small molecule antagonists of the SWI/SNF complex and TGF-β pathway inhibit SARS-CoV-2 infection *in vitro*. The hits from this screen represent novel therapeutic targets for SARS-CoV-2 and will significantly enhance our understanding of COVID-19 pathogenesis.

## Results

To identify host genes essential for cell survival in response to SARS-CoV-2 we selected the *C. sabaeus* (African green monkey) cell line Vero-E6, which is highly susceptible to SARS-CoV-2 infection and virus-induced cytopathic effects (*19–21*). We performed two independent genome-wide screens, utilizing a *C. sabaeus* genome-wide pooled CRISPR library composed of 83,963 targeting single guide RNAs (sgRNAs), with an average of four sgRNAs per gene, and 1,000 non-targeting control sgRNAs. The two screens used Vero-E6 lines expressing two different Cas9 nuclease constructs (Cas9-v1 and Cas9-v2); Cas9-v2 has an additional nuclear-localization sequence to increase activity. We transduced both Vero-Cas9 cell lines with the *C. sabaeus* sgRNA library and challenged cells with SARS-CoV-2 (**Fig. 1A**). To generate a robust dataset, we performed independent screens at different cell densities, fetal bovine serum (FBS) concentrations, and multiplicities of infection (MOI). Genomic DNA was harvested from surviving cells at 7 days post-infection (dpi) and guide abundance was determined by PCR and massively-parallel sequencing.

**Fig. 1.**
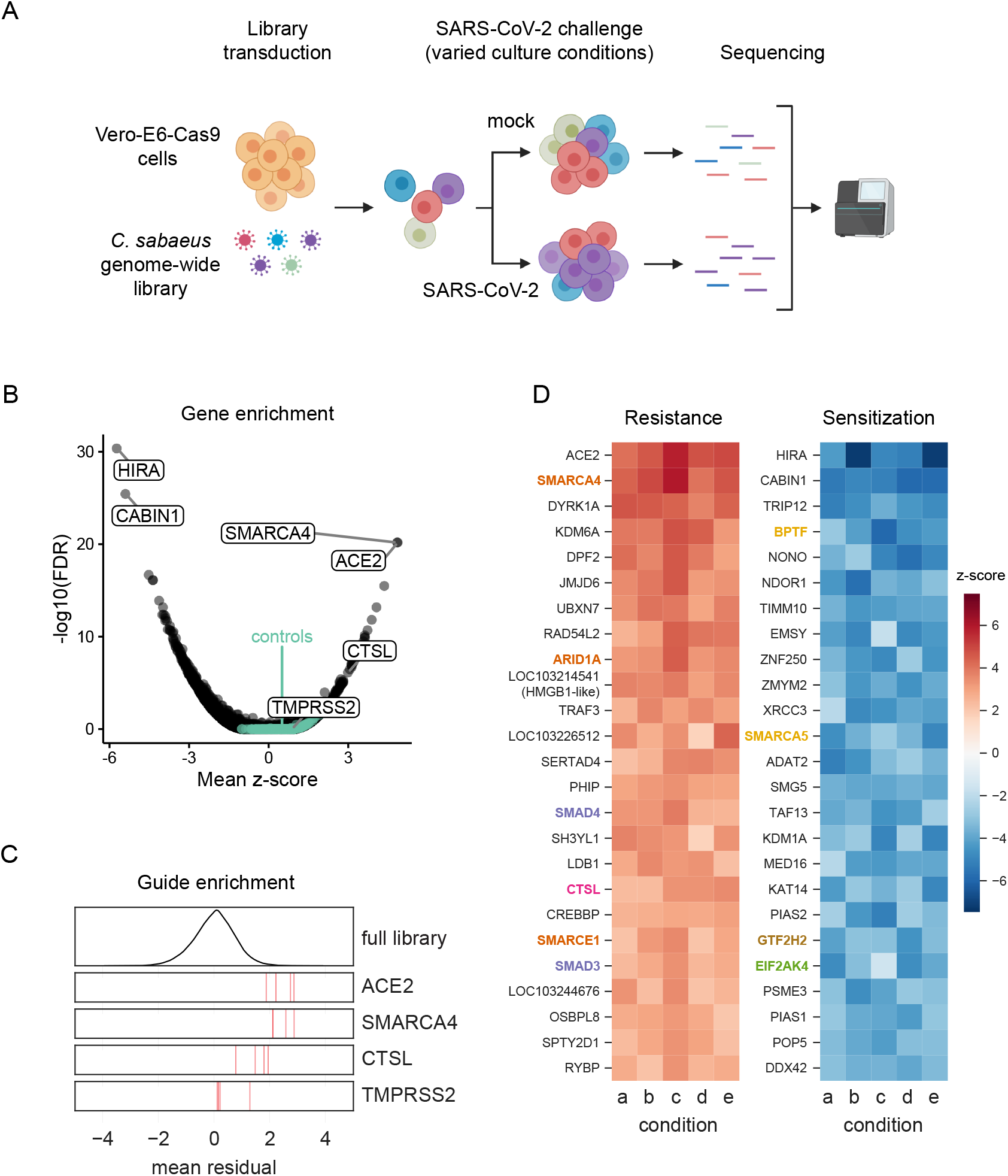
Genome-wide CRISPR screen identifies genes critical for SARS-CoV-2-induced cell death. (A) Schematic of pooled screen. Vero-E6 cells expressing Cas9 were transduced with the genome-wide C. sabaeus library via lentivirus. The transduced cell population then either received a mock treatment or was challenged with SARS-CoV-2 under various culture conditions. Surviving cells from each condition were isolated and the sgRNA sequences were amplified by PCR and sequenced. (B) Volcano plot showing top genes conferring resistance and sensitivity to SARS-CoV-2. The gene-level z-score and −log10(FDR) were both calculated using the mean of the five Cas9-v2 conditions. Non-targeting control sgRNAs were randomly grouped into sets of 4 to serve as “dummy” genes and are shown in green. (C) Performance of individual guide RNAs targeting ACE2, SMARCA4, CTSL, and TMPRSS2. The mean residual across the five Cas9-v2 conditions is plotted for the full library (top) and for the 4 guide RNAs targeting each gene. (D) Heatmaps of the top 25 gene hits for resistance and sensitivity, ranked by mean z-score in the Cas9-v2 conditions.Genes that are included in one of the gene sets labeled in (Fig 2A) are colored accordingly. Condition a: Cas9v2 D5 2.5e6 Hi-MOI; b: Cas9v2 D5 5e6 Hi-MOI; c: Cas9v2 D2 5e6 Hi-MOI; d: Cas9v2 D10 5e6 Hi-MOI; e: Cas9v2 D5 2.5e6 Lo-MOI.

To assess technical performance, we calculated the log-fold change of each guide relative to the original lentivirus plasmid pool and observed strong correlation between different cell culture conditions, with the greatest distinction between the two different Cas9 constructs (Pearson r>0.61 among Cas9-v2 conditions and r>0.46 among Cas9-v1 conditions; **Fig. S1A**). We compared the depletion of sgRNAs targeting essential genes versus sgRNAs targeting nonessential genes in the mock-infected conditions with each Cas9 construct and observed superior performance with the Cas9-v2 construct (AUC = 0.82 vs 0.70 for Cas9-v1; **Fig. S1B**), although the top positively and negatively selected hits remained concordant between the two Cas9 constructs (Pearson r = 0.24, **Fig. S1C**) (*22, 23*). The enhanced Cas9 activity of Cas9-v2 was confirmed with a GFP reporter assay (Fig. **S1D-E**). We therefore proceeded with the data from the Cas9-v2 screens and calculated a guide-level residual (representing a log2 fold change) between mock-infected and SARS-CoV-2 infected cells (**Fig. S1F**). A positive residual indicates a gene is pro-viral and knockout confers resistance to virus-induced cell death, while a negative residual indicates a gene is anti-viral and sensitizes a cell to virus-induced cell death. A z-score for enrichment or depletion was determined for each condition based on the distribution of residuals for all sgRNAs. We then averaged z-scores across all five Cas9-v2 screen conditions, combined p-values by using Fisher’s method, and calculated a false discovery rate (FDR) by using the Benjamini-Hochberg procedure to identify hit genes.

The screen recovered genes in both the resistance (pro-viral) and sensitization (anti-viral) directions, with a minimum false discovery rate of 0.030 for non-targeting controls (**Fig. 1B**), demonstrating high technical quality. The strongest resistance hit was the viral receptor *ACE2* (mean z-score = 4.9; descending rank = 1; **Fig. 1B-D**). *CTSL*, which encodes the Cathepsin L protease, was also positively selected in all conditions (mean z-score = 3.0; descending rank = 18; **Fig. 1B-D**). We did not observe enrichment of *TMPRSS2* (mean z-score = 0.9; descending rank = 2726; **Fig. 1B-C**), nor the proteases *TMPRSS4* and *FURIN* (mean z-scores = 1.1, 0.4; descending ranks = 1657, 7180, respectively), which have also been implicated in SARS-CoV-2 entry (*8, 12, 14*), suggesting these proteases are either not essential in a cell-intrinsic manner for SARS-CoV-2-induced cell death as performed herein and/or are functionally redundant. We also observed minimal overlap with a recent SARS-CoV-2 protein interactome conducted in HEK293T cells (**Fig. S2**), indicating that our functional genomic screen identified novel host genes regulating SARS-CoV-2 infection (*7*).

To systematically identify hit gene sets, we used the STRING-db enrichment detection tool and identified 623 significant gene sets (**Fig. S3**) from 10 sources (e.g. KEGG, GO process) (*24*). The top gene sets that scored in the positive direction (pro-viral), negative direction (ant-viral), or both directions are shown in **Fig. 2A** and their respective genes are shown in **Fig. 2B-H**. *SMARCA4* (*BRG1*), the catalytic subunit of the SWI/SNF remodeling complex, scored after *ACE2* as the second-strongest resistance hit (mean z-score = 4.9; descending rank = 2; **Fig. 1B-D**), with several other members including *ARID1A, SMARCE1, SMARCB1*, and *SMARCC1* showing enrichment (z-scores = 3.6, 2.8, 2.4, 2.3, descending ranks = 9, 20, 47, 59, respectively; **Fig. 2B**). The SWI/SNF complex is an ATP-dependent nucleosome remodeling complex that regulates chromatin accessibility and gene expression (*25, 26*). Similarly, we identified several other histone modifying enzymes as key regulators of SARS-CoV-2-induced cell death (**Fig. 1D**). Other pro-viral genes include the histone demethylase *KDM6A* (z-score = 4.1; descending rank = 4), histone methyltransferase *KMT2D* (z-score = 2.6; descending rank = 35), as well as the lysyl hydroxylase *JMJD6* (z-score = 3.7; descending rank = 6) (**Fig. 1D**) (*27–30*). In contrast, sgRNAs targeting *HIRA, CABIN1*, and *ASF1A* were negatively selected revealing an anti-viral function. These genes represent three of the four components of the HUCA histone H3.3 chaperone complex (z-scores = −5.7, −5.4, −3.0; ascending ranks = 1, 2, 64, respectively), suggesting an anti-viral role for the deposition of the histone variant H3.3 (*31*).

**Fig. 2.**
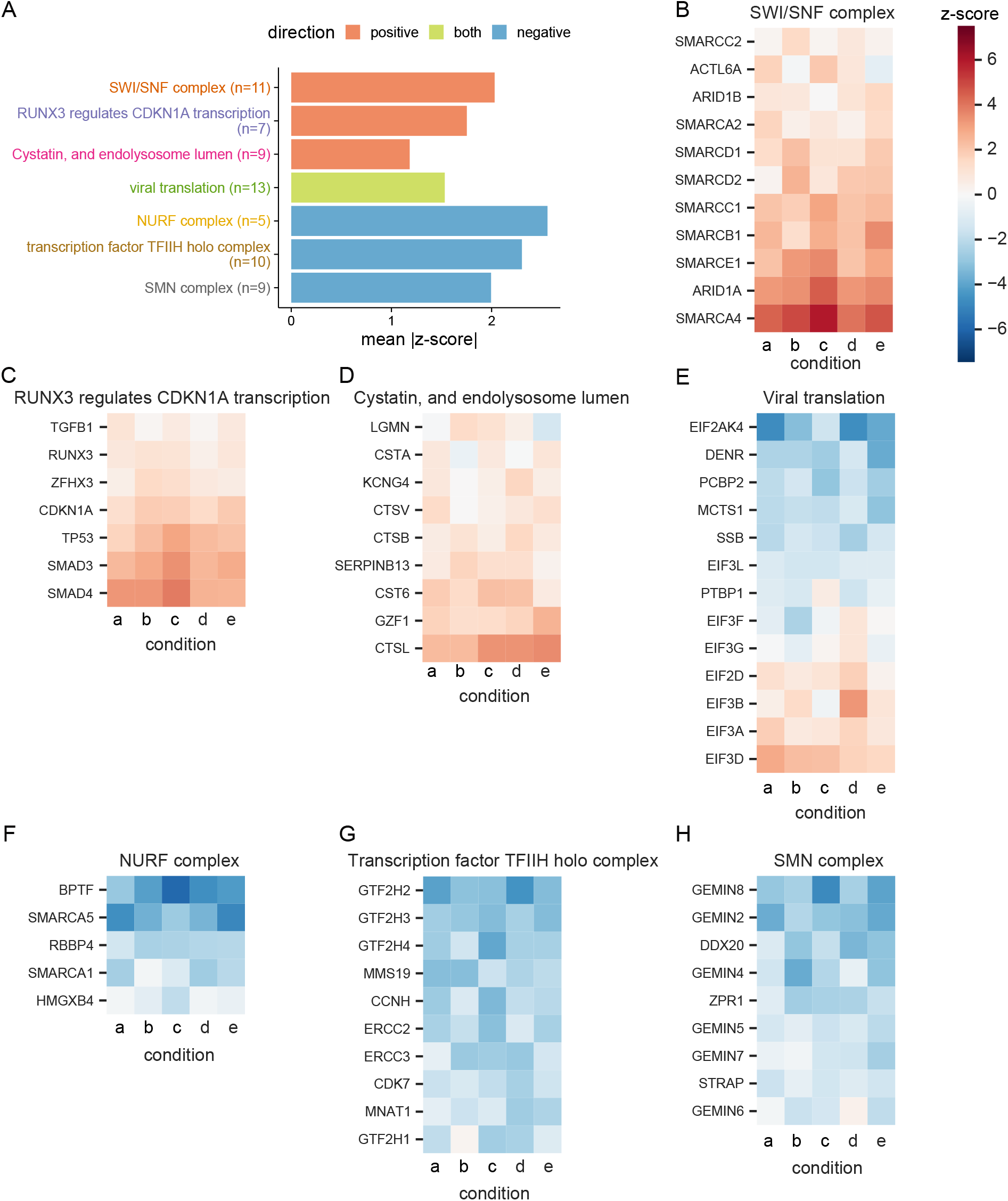
Performance of genes in top selected gene sets. (A) Top three gene sets, which score in the positive direction (resistance), negative direction (sensitization), or both, filtered for gene sets with at least five genes and which are most central to a given module (Fig. S3) and then ranked by mean absolute z-score. There was only one gene set which met these criteria and scored on both ends. The number of genes in each set is indicated in parentheses. For each gene in the “SWI/SNF complex” gene set from STRING, the z-score in each culture condition is shown. Similarly, the genes in the gene sets (B) “RUNX3 regulates CDKN1A transcription” from Reactome, (C) “Cystatin, and endolysosome lumen” from STRING, (D) “viral translation” from GO, (E) “NURF complex” from GO, (F) “transcription factor TFIIH holo complex” from GO, and (G) “SMN complex” from GO are shown. Condition a: Cas9v2 D5 2.5e6 Hi-MOI; b: Cas9v2 D5 5e6 Hi-MOI; c: Cas9v2 D2 5e6 Hi-MOI; d: Cas9v2 D10 5e6 Hi-MOI; e: Cas9v2 D5 2.5e6 Lo-MOI.

We additionally observed enrichment in the *“RUNX3* regulates *CDKN1A* transcription” gene set from Reactome (**Fig. 2A, 2C**). RUNX3 binds to the transcription factors SMAD3 and SMAD4 and induces *CDKN1A* expression in response to TGF-β (*32, 33*). The TGF-β signaling pathway is involved in diverse physiologic processes including cell proliferation, differentiation, and apoptosis (*34–36*). Ligand binding to a TGF-β superfamily receptor causes phosphorylation of SMAD transcription factors in the cytoplasm and results in nuclear translocation and target gene transcription (*37, 38*). In addition to observing enrichment of sgRNAs targeting the signal transducers *SMAD3* and *SMAD4* (z-scores = 2.8, 3.1; descending ranks = 21, 15, respectively), we identified the TGF-β superfamily receptor *ACVR1B* (z-score = 2.2; descending rank = 65) as pro-viral. Genes encoding the TGF-β cytokine, other TGF-β superfamily receptors, and additional SMADs were not enriched, suggesting specificity of the *ACVR1B/SMAD3/SMAD4* axis in SARS-CoV-2-induced cell death. The “cystatin and endolysosomal lumen” gene set, which includes *CTSL*, was one of the top gene sets enriched in the CRISPR screen (**Fig. 2D**). The set “viral translation” had genes enrich in both the positive and negative sides of the screen (**Fig. 2E**), while the top three anti-viral gene sets were the NURF complex, TFIIH holo complex and SMN complex (**Fig. 2F-H**).

We selected 25 genes for further validation, consisting of 18 resistance and 7 sensitization genes (**Fig. 3A**). We transduced Vero-Cas9 cells with one of 42 individual sgRNAs (1 to 3 sgRNAs per gene), challenged each of the 42 cell lines with SARS-CoV-2, and assessed cell viability at 3 dpi. Cells receiving sgRNAs targeting pro-viral genes exhibited greater viability than those with non-targeting control sgRNAs, while all genes identified as antiviral in the primary screen exhibited increased susceptibility to cell death relative to controls, confirming the efficiency and reproducibility of the screen (**Fig. 3B-C**).

**Fig. 3.**
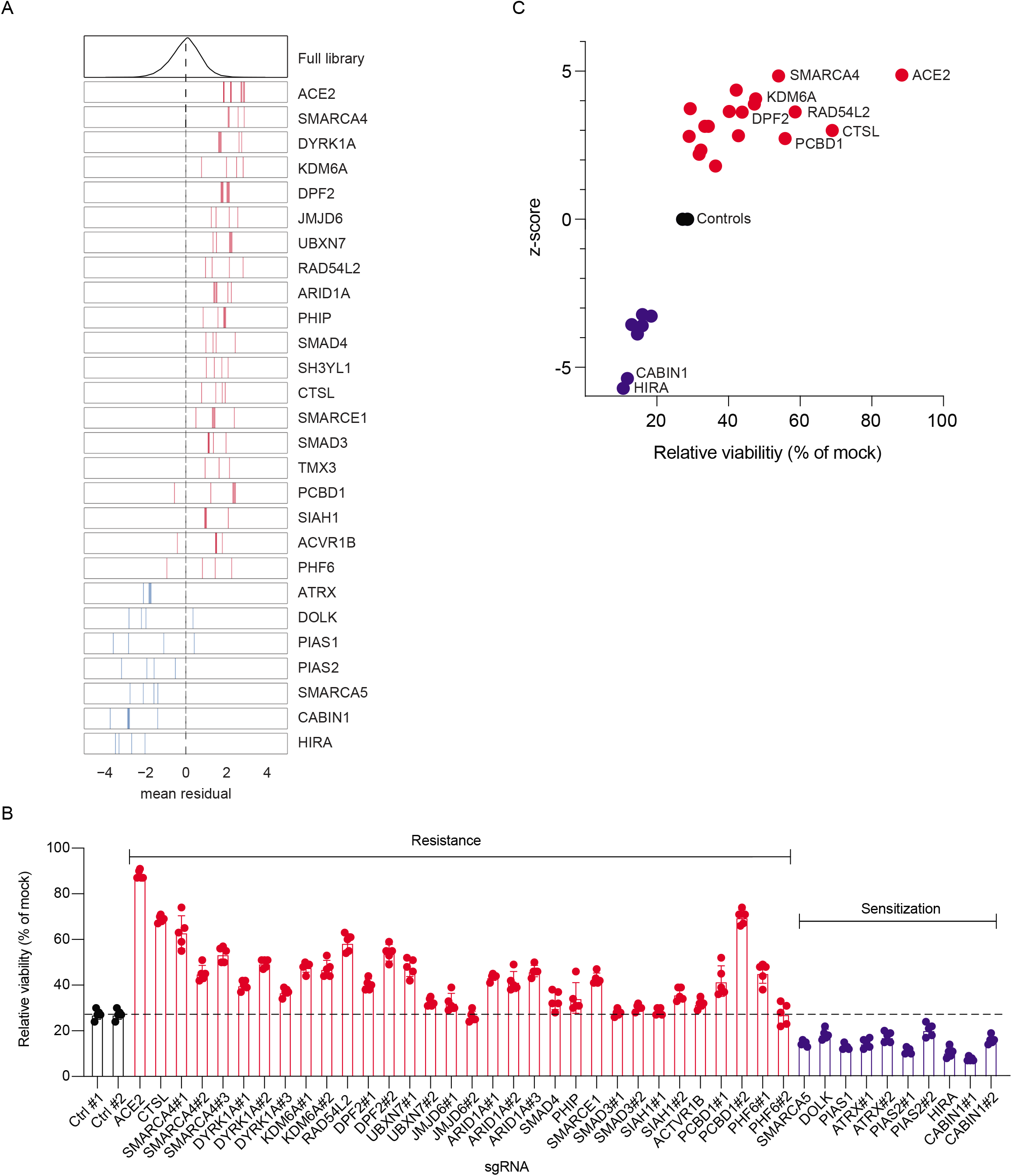
Arrayed validation of 18 resistance and 7 sensitization hit genes. (A) Performance in the pooled screen of guide RNAs targeting the 25 genes selected for further validation. The mean residual across the five Cas9-v2 conditions is plotted for the full library (top) and for the 3-4 guide RNAs targeting each gene. Genes that scored as resistance hits are shown in red; genes that scored as sensitization hits are shown in blue. The dashed line indicates a residual of 0. (B) 42 unique sgRNAs targeting 25 genes were introduced into VeroE6 cells. SARS-CoV-2 was added at MOI 0.2 and cell viability was measured at 3 dpi. (C). Z-scores from CRISPR screen correlate with cell viability of individually disrupted genes. Genes with multiple sgRNAs from (B) are averaged to generate one point per gene in (C). Data in (B) and (C) is representative of one of three independent experiments each done in quintuplicate.

The determinants of SARS-CoV-2 cell tropism are incompletely understood. We recently performed single-cell RNA sequencing of experimentally infected primary human bronchial epithelial cells. Eight epithelial cell populations were observed and ciliated cells, basal cells, and club cells were the primary target cells of infection (*39*). In the single-cell transcriptomic dataset, *ACE2, CTSL*, and *TMPRSS2* expressions were of limited value in predicting whether a cell was infected. To evaluate whether SARS-CoV-2-interacting genes identified in the CRISPR screen were predictive of infection, we assessed the predictive value of the top 50 resistance and sensitization genes for the ability to predict SARS-CoV-2 infection of primary human bronchial epithelial cells (*39*). We found that these 100 genes obtained from the CRISPR screen are significantly predictive (Wilcoxon rank sum, see Methods) of viral infection, suggesting that these genes are involved in variability of infection at the single cell level. Surprisingly, *AP2M1* and not *ACE2* was the most predictive gene in this dataset (**Fig. S4**). AP2M1 was also one of only three genes enriched in both the CRISPR screen SARS-CoV-2 protein interactome (**Fig. S2**)

The screen also revealed a putative pro-viral role for the gene *LOC103214541*, which is annotated as “HMGB1-like” in the *C. sabaeus* genome (z-score = 3.6, descending rank = 10; **Fig. 4A**). HMGB1 is a nuclear protein that binds DNA but translocates to the cytoplasm under conditions of stress, and can be secreted extracellularly where it functions as an alarmin (*40*). To validate the role of HMGB1 in SARS-CoV-2 infection, we introduced three independent sgRNAs targeting *HMGB1* into Vero-E6 cells. We observed substantial depletion of HMGB1 protein in two of the three cell lines as measured by western blot (**Fig. 4B**). *HMGB1* knockout protected cells from SARS-CoV-2-induced cell death, and the degree of protection tracked with HMGB1 levels (**Fig. 4C**). We then performed SARS-CoV-2 growth curves on control and *HMGB1* disrupted cells and observed an approximately 2-log reduction in SARS-CoV-2 replication at both 24 and 48 hours post-infection (**Fig. 4D**). To determine whether HMGB1 acted at the level of viral entry or post-entry, we pseudotyped the SARS-CoV-2 spike onto a replication defective vesicular stomatitis virus (VSV) core expressing *Renilla* luciferase (SARS2-VSVpp) and used pseudovirus expressing the VSV glycoprotein G as a control (G-VSVpp) (*41*). SARS2-VSVpp and G-VSVpp similarly infected both *HMGB1* knockout and control cells, demonstrating that HMGB1 acts after viral entry (**Fig. 4E**).

**Fig. 4.**
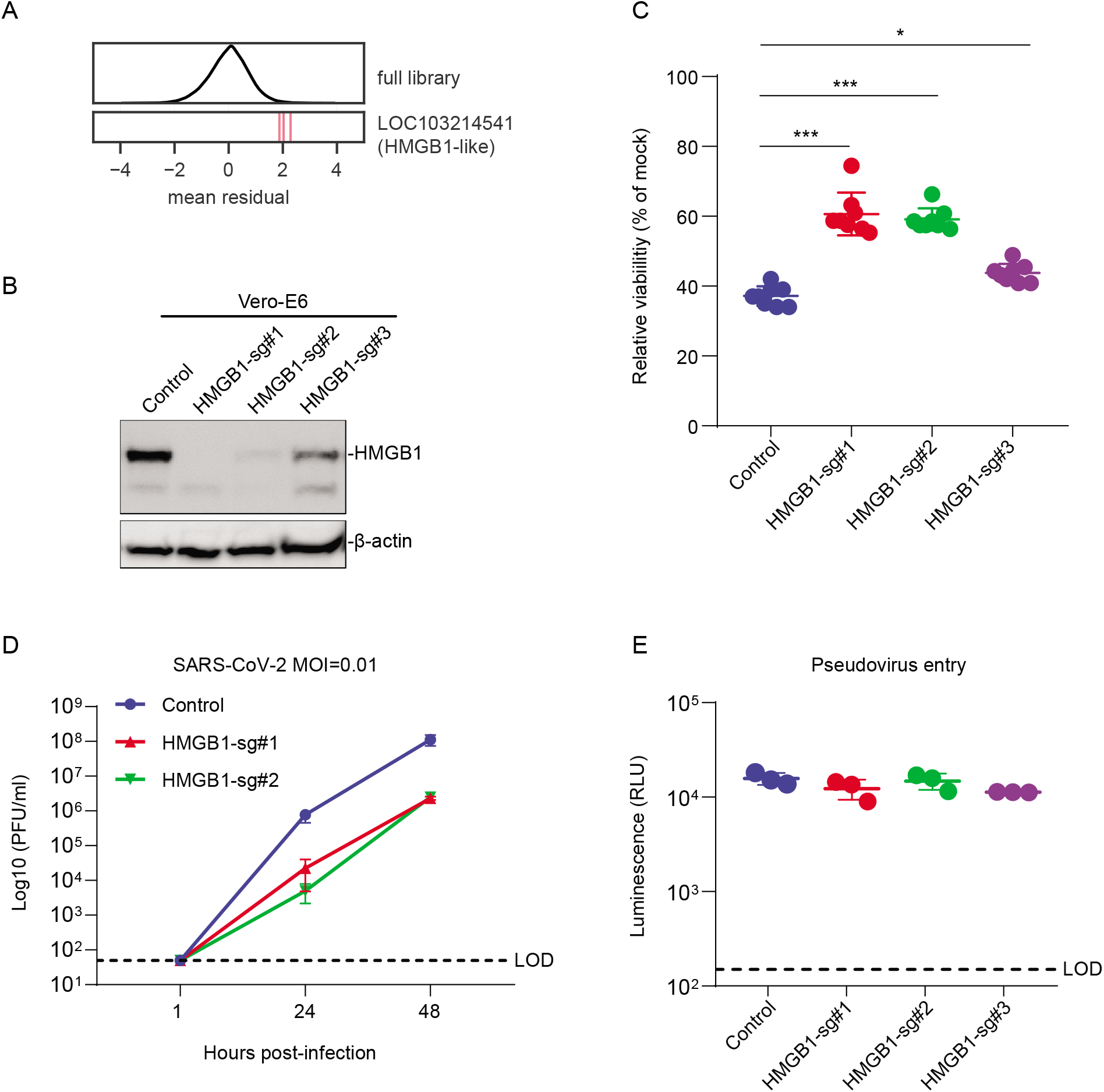
HMGB1 is required for efficient SARS-CoV-2 induced cell death. (A) Performance of individual guide RNAs targeting LOC103214541 (HMGB1-like). The mean residual across the five Cas9-v2 conditions is plotted for the full library (top) and for the 3 guide RNAs targeting that gene. (B) Western blot for HMGB1 expression in control and HMGB1 sgRNAs transduced Vero-E6 cells. (C) Control and HMGB1 sgRNAs transduced Vero-E6 cells were infected with SARS-CoV-2 at a MOI of 0.2. Cell viability relative to an uninfected control was measured 3dpi with CellTiter Glo. (D) Vero-E6 cells were infected with SARS-CoV-2 at a MOI of 0.1. Viral production as measured by plaque forming units (PFU/ml) was determined by plaque assay. (E) Vero-E6 cells were infected with VSV-G or SARS-CoV-2-Spike pseudovirus. Luminescence was measured 1dpi. LOD, limit of detection. Data was analyzed by Mann-Whitney test. Shown are means ± SEM. ns, not statistically significant; *P < 0.05; **P < 0.01; ***P < 0.001. Data in (C) are one representative experiment from three independent times each in octuplicate. Data in (D-E) are one representative experiment from two independent experiments each performed in triplicate.

Next, we asked whether the genes and pathways revealed by the screen could be targeted therapeutically. We selected small molecule antagonists previously described to inhibit these gene products and investigated their effects on SARS-CoV-2 induced cell death. In addition, we utilized a replication competent infectious clone of SARS-CoV-2 expressing the fluorescent reporter mNeonGreen (icSARS-CoV-2-mNG) to quantify viral replication in the presence of drugs (*42*). We observed dose-dependent inhibition of SARS-CoV-2 induced cell death and virus replication with Calpain Inhibitor III, which targets Cathepsin L (**Fig. 5A, E-F**) (*43, 44*). Surprisingly, we observed strong cell toxicity of the PIKFyve kinase inhibitor APY0201 even at 0.6 μM concentration. However, we did not observe a dose-dependent protective effect with APY0201, which was recently described to protect from SARS-CoV-2-induced cell death (**Fig. 5B**) (*45*), nor did we observe enrichment of PIKFyve in the CRISPR screen (z-score = −1.14). Given the pro-viral role of genes involved in TGF-β signaling, the SWI/SNF complex, and histone regulation, we tested whether existing small molecule antagonists of these pathways have antiviral activity against SARS-CoV-2. Specifically, treatment with PFI-3, which inhibits the bromodomains of the SWI/SNF proteins SMARCA4 and SMARCA2 (*42, 46*), partially protected cells from virus-induced death (**Fig. 5C**) and reduced frequency of viral infection, as measured by expression of mNeonGreen (**Fig. 5E-F**). We also assessed the TGF-β pathway with the small molecule SIS3, an inhibitor of SMAD3, which was enriched in the genetic screen (*47*). SIS3 exhibited dosedependent protection from virus-induced cell death and also inhibited SARS-CoV-2 replication (**Fig. 5D-F**). This provides pharmacological validation in addition to genetic evidence that these pathways are critical to SARS-CoV-2 infection *in vitro*.

**Fig. 5.**
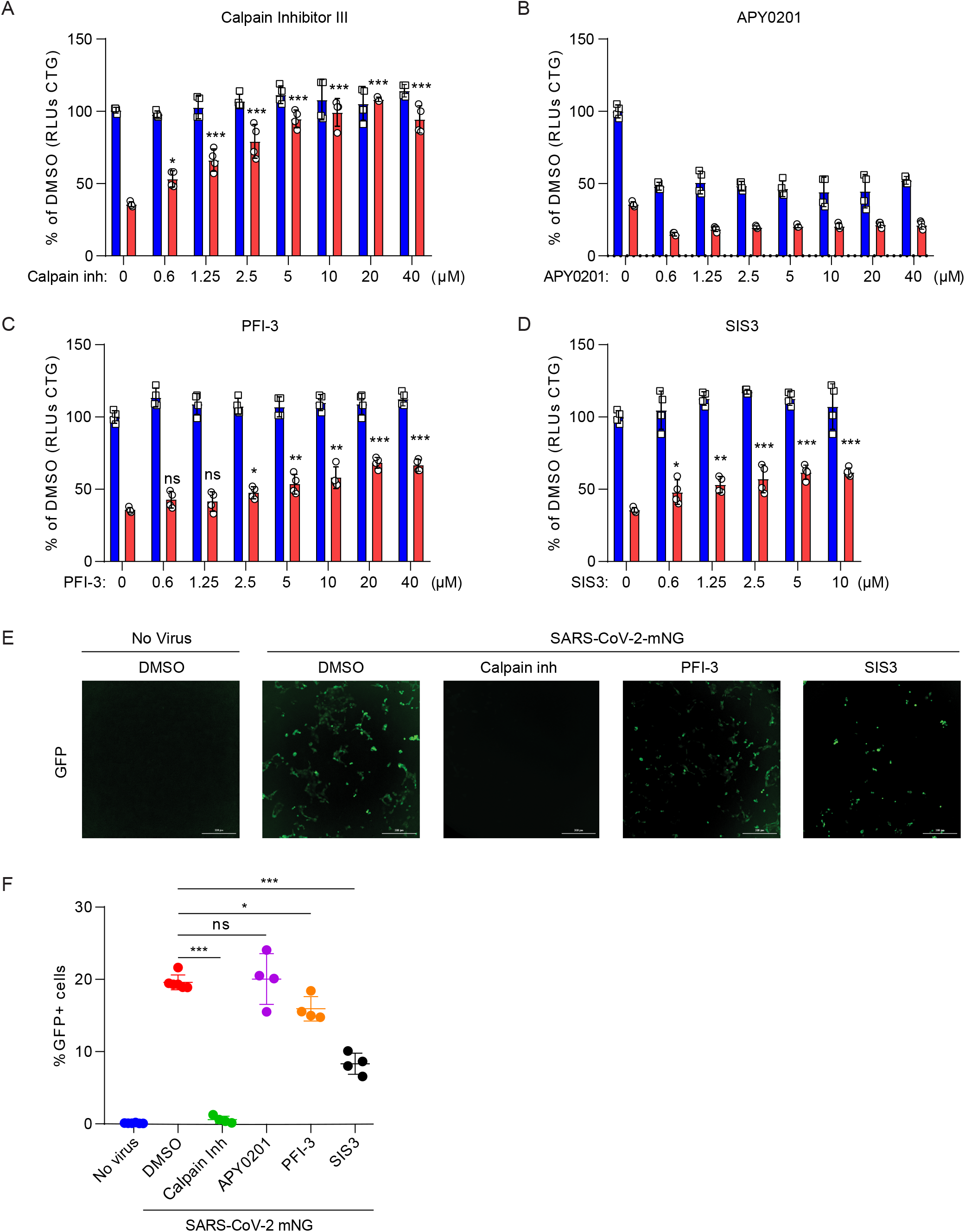
Small molecules protect cells from SARS-CoV-2 induced cell death. (A to D) Vero-E6 cells were pretreated with the indicated concentration of Cathepsin L inhibitor Calpain Inhibitor III (A), PIKfyve inhibitor APY0201 (B) SMARCA4 inhibitor PFI-3 (C), SMAD3 inhibitor SIS3 (D) for 48 hours and then infected with SARS-CoV-2 at a MOI of 0.2. Cell viability was measured at 3dpi and compared to mock infected controls. Red, infected; blue, mock (E to F) Vero-E6 cells were pretreated with 10 μM Calpain inhibitor III, PFI-3 or SIS3 for 48 hours and then infected with icSARS-CoV-2 mNG at a MOI of 1. Infected cell frequencies were measured by mNeonGreen expression at 2 dpi. Data were analyzed by one-way ANOVA with Tukey’s multiple comparison test. Shown are means ± SEM. ns, not statistically significant; *P < 0.05; **P < 0.01; ***P < 0.001. Data are pooled from two independent experiments each in duplicate except in (E), where results are shown from one representative experiment.

## Discussion

We generated a novel CRISPR library and performed the first genome-wide CRISPR screen with SARS-CoV-2. The identification of known pro-viral genes *ACE2* and *CTSL* demonstrate the technical quality of the screen, providing confidence in the additional genes that regulate SARS-CoV-2 infection (*8, 14, 15, 44*). We discovered genes involved in diverse biological processes including chromatin remodeling, histone modification, cellular signaling, and RNA regulation. We assessed 25 of these genes with individual sgRNAs in an arrayed format, including both pro-viral and antiviral genes, and identified small molecule antagonists that confer protection to SARS-CoV-2-induced cell death and infection. Importantly, this study is not without limitations as we used Vero-E6 cells, a type I interferon-deficient non-human primate cell line, rather than primary human cells (*48–50*). However, Vero-E6 cells are a model cell line for isolating viruses and were selected based on their susceptibility to SARS-CoV-2-induced death. Further, African green monkeys are susceptible to SARS-CoV-2 infection (*19–21*). Interestingly, we did not detect several genes previously implicated in SARS-CoV-2 infection including *TMPRSS2, PIKFyve*, and *TPC2* suggesting that these genes are either not essential in a Vero cell-intrinsic manner or that functional redundancies exist with other genes (*15, 45*). Future work is necessary to evaluate the genes identified here in human cells and animal models.

The abundance of genes identified here with functional roles in chromatin regulation and histone modification were surprising, as SARS-CoV-2 is an RNA virus that replicates in the cytoplasm. However, these genes highlight the potential importance of epigenetic regulation of SARS-CoV-2 infection and pathogenesis. Epigenetic processes have previously been implicated in regulating antigen presentation and interferon-stimulated gene induction after MERS-CoV and SARS-CoV infection (*51–53*); however, given that Vero-E6 cells are type I interferon-deficient, distinct mechanism(s) may be at play. Here we identify a novel role for HMGB1 in SARS-CoV-2 susceptibility. HMGB1 is a predominantly nuclear yet pleiotropic protein. It binds nucleosomes regulating chromatin in the nucleus, acts as a sentinel of non-self nucleic acids, transports genetic material, and functions as a secreted alarmin in response to virus infection (*53–55*). Interestingly, anti-HMGB1 therapies can reduce respiratory syncytial virus replication and influenza A-induced lung pathology in animal models (*56, 57*), while in adenovirus infection, adenovirus protein VII binds HMGB1 and inhibits its pro-inflammatory functions (*58*). Elucidating the molecular interactions between SARS-CoV-2 and HMGB1 is an important area of future research.

The SWI/SNF chromatin remodeling complex and components of the TGF-β pathway were both identified as pro-viral pathways. It is intriguing to speculate that the TGF-β signaling pathway, through Activin 1RB, SMAD3, and SMAD4, may promote the SWI/SNF-driven changes to chromatin architecture, thus inducing expression of a pro-viral gene expression program. Interestingly, the nucleoprotein of SARS-CoV was shown to bind SMAD3, preventing the SMAD3/SMAD4 interactions that drive downstream of TGF-β signaling and virus-induced apoptosis (*59*). Further, *SMARCA4*, which encodes the catalytic component of the SWI/SNF complex, can interact with SMAD3 and is critical for induction of many TGF-β responsive genes (*60*). We speculate that this gene-expression program may be inhibited by the histone H3.3 (HIRA chaperone) complex, which both co-precipitates with and has overlapping chromatin binding sites as SMARCA4 (*61*). The histone H3.3 complex can also be induced by TGF-β (*62*). Finally, Cathepsin L can cleave histone H3 tails, which in turn regulate cellular senescence (*59, 63*). Whether this process is manipulated by SARS-CoV-2 infection remains an area for future investigation.

The pro-viral and antiviral genes identified herein have important implications for our understanding of COVID-19 pathogenesis, therapeutics, and vaccine design. First, SARS-CoV-2 can cause diverse phenotypes ranging from asymptomatic infection to severe respiratory failure and death (*64, 65*). The basis for this variation among people and between species is unclear. The genes and pathways identified here may explain this variation as disease susceptibility may positively correlate with expression of resistance genes and negatively correlate with sensitization genes on the cellular, tissue, and organismal level. For example, cigarette smoking both increases *ACE2* expression and exacerbates COVID-19 pathogenesis (*66*). The regulatory network underlying this is unknown, but it is intriguing to speculate that the chromatin and histone modifying genes identified here contribute to expression of a heterogeneous pro-viral gene expression program that potentially regulates *ACE2* and other viral interacting genes. Interestingly, the second-ranked hit in our screen, SMARCA4, was previously demonstrated to regulate *ACE2* expression under conditions of stress in cardiac endothelial cells (*67*). SMARCA4 has also been shown to promote proliferation of alveolar epithelial cells, a major SARS-CoV-2 target, and protect against pulmonary fibrosis, a common sequelae of both SARS-CoV and SARS-CoV-2 (*68–70*).

The genetic screen revealed novel therapeutic targets for SARS-CoV-2 infection. As a proof-of-principle, we tested several small molecule inhibitors and identified three molecules that inhibit SARS-CoV-2 replication and virus-induced cell death. Therapeutic targeting of these genes and pathways, including the SWI/SNF complex and TGF-β pathway, may prove clinically useful. Additionally, we identified and validated anti-viral genes, including regulators of the histone variant H3.3 (*CABIN1, HIRA, ASF1A) (31, 71*). While these genes potentially provide protection from SARS-CoV-2, they may also prove fruitful in generating knockout cell lines with increased susceptibility to diverse human coronaviruses, which may facilitate coronavirus vaccine production.

SARS-CoV, MERS-CoV, and SARS-CoV-2 reveal the pandemic potential and dangers of emerging coronaviruses for which there are no FDA-approved therapeutics or vaccines (*5–7*). To our knowledge this study represents the first genome-wide genetic screen performed with any coronavirus. Ultimately, our findings may be broadly applicable to other human and emerging coronaviruses which may facilitate development of host-directed therapies against existing and future pandemic coronaviruses.

## Acknowledgements

We would like to acknowledge Brian Smith, Nancy Brown, and Ruth Montgomery for generous resources; Douglas E. Brackney and Kenneth Plante for critical reagents; Marissa Feeley and Yenarae Lee for technical assistance; Stephanie Eisenbarth, David Schatz, Richard Flavell, Scott Pope, and Ruslan Medzhitov for helpful discussions; Benjamin Fontes and Daniele Scavone for Environmental Health and Safety.

## Funding

This work was supported by NIH grants K08 AI128043 (CBW), U19 AI133524 (JGD), P50 CA121974 (QY), R01 AI087925 (BDL), T32GM007223 (WJLC), T32GM007205 (WJLC, JSC), F30HL149151 (JSC), R01 AI123449 (BL), F31 AI54739 (KO), Burroughs Wellcome Fund Career Award for Medical Scientists (CBW), Ludwig Family Foundation (CBW), and Emergent Ventures Fast Grant (CBW, DvD), Merkin Institute Fellowship (JGD).

## Author contributions

**Jin Wei**: Conceptualization, Methodology, Validation, Formal Analysis, Investigation, Writing - Original Draft, Visualization, **Mia Madel Alfajaro:** Conceptualization, Methodology, Validation, Formal Analysis, Investigation, **Ruth E. Hanna:** Methodology, Formal Analysis, Data Curation, Writing - Original Draft, Visualization, **Peter C. DeWeirdt:** Writing - Original Draft, Software, Formal Analysis, Data Curation, Visualization, **Madison S. Strine:** Validation, Formal Analysis, Investigation, **William J. Lu-Culligan:** Validation, Investigation, **Shang-Min Zhang:** Validation, **Laura Abriola**: Investigation, **Yulia Surovtseva:** Investigation, **Cameron O. Schmitz:** Validation, Investigation, **Jennifer S. Chen**: Validation, Investigation, **Vincent R. Graziano:** Validation, Investigation, **Madeleine Mankowski:** Validation, Investigation, **Renata B. Filler:** Investigation, **Victor Gasques:** Methodology, Formal Analysis, **Fernando de Miguel:** Validation, **Huacui Chen**: Validation, **Kasopefoluwa Oguntuyo:** Resources, **Robert C. Orchard:** Conceptualization, Methodology, **Benhur Lee:** Resources, Supervision, **Brett Lindenbach:** Resources, Methodology, **Katerina Politi:** Resources, Supervision, **David van Dijk:** Resources, Supervision, Methodology, Formal Analysis **Matthew D. Simon:** Resources, Supervision, Conceptualization, Methodology, **Qin Yan:** Resources, Supervision, Conceptualization, Methodology, **John G. Doench:** Conceptualization, Methodology, Formal Analysis, Resources, Writing - Original Draft, Supervision, **Craig B. Wilen:** Conceptualization, Methodology, Investigation, Resources, Writing - Original Draft, Supervision, Funding Acquisition. All authors reviewed and edited the manuscript.

## Competing interests

Yale University (CBW) has a patent pending related to this work entitled: “Compounds and Compositions for Treating, Ameliorating, and/or Preventing SARS-CoV-2 Infection and/or Complications Thereof.” Yale University has committed to rapidly executable non-exclusive royalty-free licenses to intellectual property rights for the purpose of making and distributing products to prevent, diagnose and treat COVID-19 infection during the pandemic and for a short period thereafter. JGD consults for Foghorn Therapeutics, Maze Therapeutics, Merck, Agios, and Pfizer; JGD consults for and has equity in Tango Therapeutics.

## Material availability

Materials will be made publicly available either through publicly available repositories or via the authors at the time of publication of a peer-reviewed manuscript upon execution of a Material Transfer Agreement.

## Materials and Methods

### Cell culture

Vero-E6 cells were cultured in Dulbecco’s Modified Eagle Medium (DMEM) with 10% heat-inactivated fetal bovine serum (FBS), and 1% Penicillin/Streptomycin unless otherwise indicated. For Vero-E6 cells, 5 μg/ml of puromycin (Gibco) and 5 μg/ml blasticidin (Gibco), were added as appropriate.

### Generation of SARS-CoV-2 stocks

To generate SARS-CoV-2 viral stocks, Huh7.5 cells were inoculated with SARS-CoV-2 isolate USA-WA1/2020 (BEI Resources #NR-52281) at an MOI of approximately 0.01 for three days to generate a P1 stock. The P1 stock was then used to inoculate Vero-E6 cells for three days at approximately 50% cytopathic effects. Supernatant was harvested and clarified by centrifugation (450 *g* x 5 min) and filtered through a 0.45-micron filter, and then aliquoted for storage at −80°C. Virus titer was determined by plaque assay using Vero-E6 cells. To generate icSARS-CoV-2-mNG stocks, lyophilized icSARS-CoV-2-mNG was resuspended in 0.5 ml of deionized water and then 50 μl of virus was diluted in 5 ml media (*42*). icSARS-CoV-2-mNG was provided by the World Reference Center for Emerging Viruses and Arboviruses (Galveston, TX). This was then added to 10^7^ Vero-E6 cells in a T175 flask. At 3 dpi, the supernatant was collected and clarified by centrifugation (450 *g* x 5 min), filtered through a 0.45-micron filter, and aliquoted for storage at - 80°C. All work with infectious virus was performed in a Biosafety Level 3 laboratory and approved by the Yale University Biosafety Committee.

### SARS-CoV-2 plaque assays

Vero-E6 cells were seeded at 7.5 x 10^5^ cells/well in 6-well plates or 4 x 10^5^ cells/well in 12-well plates. The following day, media were removed and replaced with 100 μl of 10-fold serial dilutions of virus, and plates were incubated at 37°C for 1 hour with gentle rocking. Subsequently, overlay media (DMEM, 2% FBS, 0.6% Avicel RC-581) was added to each well. At 2 dpi, plates were fixed with 10% formaldehyde for 30 min and then stained with crystal violet solution (0.5% crystal violet in 20% ethanol) for 30 min, then rinsed with deionized water to visualize plaques.

### CRISPR screen

Vero-E6 cells (ATCC) were transduced with lenti-Cas9 (Cas9-v1, Addgene 52962) or pLX_311-Cas9 (Cas9-v2, Addgene 96924) and selected with blasticidin (5 μg/ml) for 10 days. Cas9 activity was assessed by transducing parental Vero-E6 or Vero-E6-Cas9 cells with pXPR_047 (Addgene 107645), which expresses eGFP and an sgRNA targeting eGFP (*72*). Cells were transduced for 24 hour, selected for five days with puromycin, and the frequency of eGFP expression was assessed by flow cytometry on a Cytoflex S (Beckman). The African green monkey (AGM) genome-wide CRISPR knockout library, which contains four unique sgRNA per gene in pXPR_050 (Addgene 96925) and was designed according to the same general principles as the ‘Brunello’ human genome-wide library (*74*) was delivered by lentiviral transduction of 2 x 10^8^ Vero-E6-Cas9 at ~0.3 MOI. This equates to 6 x 10^7^ transduced cells, which is sufficient for the integration of each sgRNA into ~750 unique cells. Two days post-transduction, puromycin was added to the media and transduced cells were selected for seven days. Two infection conditions were set up for the screening with Cas9-v1: (1) 10% FBS, 5 x 10^6^ cells, MOI 0.1 (2) 10% FBS, 5 x 10^6^ cells, MOI 0.01. Five infection conditions were set up for the screening with Cas9-v2: (1) 10% FBS, 5 x 10^6^ cells, MOI 0.1; (2) 5% FBS, 5 x 10^6^ cells, MOI 0.1; (3) 5% FBS, 2.5 x 10^6^ cells, MOI 0.1; (4) 5% FBS, 2.5 x 10^6^ cells, MOI 0.01; (5) 2% FBS, 5 x 10^6^ cells, MOI 0.1. For each condition, 4 x 10^7^ cells were seeded in T175 flasks at the indicated cell concentration. For the Cas9-v1 screen, the mock sample was plated in 10% FBS at 5 x 10^6^ cells in each of eight T175 flasks. For the Cas9-v2 screen, the mock sample was plated identically to condition (2) above in 5% FBS at 5 x 10^6^ cells in each of eight T175 flasks. Cells were infected with SARS-CoV-2 at the indicated MOI. Mock infected cells were harvested 48 hours after seeding and served as a reference for sgRNA enrichment analysis. At 7 dpi, cell lysates were harvested in DNA/RNA shield (Zymo Research) and genomic DNA (gDNA) of surviving cells was isolated using a genomic DNA cleanup kit according to manufacturer instructions (Zymo Research, D4065). For Illumina sequencing and screening analysis, PCR was performed on gDNA to construct Illumina sequencing libraries, with each well containing 10 μg gDNA as described previously (*73, 74*). The lentiviral plasmid DNA pool was also sequenced as a reference.

### Screen analysis

Guide sequences were extracted from the sequencing reads with PoolQ version 3.2.9 (Broad Institute, https://portals.broadinstitute.org/gpp/public/software/poolq), using a “CACCG” search prefix, and a counts matrix was generated. Read counts were log-normalized within each condition using the following formula:

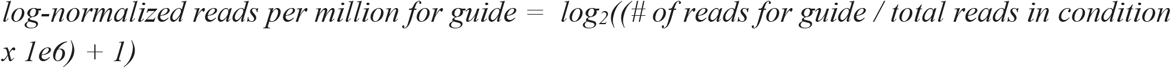

Prior to analysis, any sgRNAs with an outlier abundance in the plasmid DNA pool (defined as a log-normalized read count > 3 standard deviations from the mean) or that had > 5 predicted off-target sites with a CFD score = 1 (“Match Bin I”) were filtered out. This removed 755 sgRNAs; the remaining 84,208 sgRNAs were used for all analyses. We then calculated log-fold changes (LFCs) by subtracting the log-normalized plasmid DNA. For each condition, we fit a natural cubic spline with 4 degrees of freedom, using the mock infected LFCs as the independent variable and the relevant condition’s LFCs as the dependent variable. We used the residual from this fit spline to represent the deviation from the expected LFC for each guide. To combine these residuals at the gene level, we calculated a z-score for each condition, 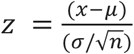, where *x* is the mean residual for a gene, *μ*is the mean residual of all sgRNAs, σis the standard deviation of all sgRNAs and nis the number of sgRNAs for a given gene. We used the normal distribution function to calculate p-values from the z-scores. To combine p-values across multiple conditions we used Fisher’s method. Finally, to calculate the false discovery rate for each gene we used the Benjamini-Hochberg procedure.

### Gene set enrichment and network analysis

We used the STRING enrichment detection tool to identify significantly enriched gene sets, using African green monkey gene symbols, but testing for enrichment across human gene sets. We analyzed sets from all available sources provided by that tool, including sets of clusters of proteinprotein interactors in STRING, and excluding the PubMed gene sets. We then generated a network of enriched gene sets by drawing edges between sets with a significant overlap between genes. We evaluated the significance of overlap using Fisher’s exact test. We clustered the network with the infomap algorithm (*75*), using the fraction of genes which overlap between any two sets as edge weights, and the mean absolute z-score as node weights. We evaluated the centrality of each node to a given cluster using the PageRank algorithm with a damping factor of 0.5 and employing the same edge and node weights (for personalized PageRank) as we did with clustering (*76*).

### Cell infection prediction model

An XGBoost model was trained to predict infected cells based on 24,714 genes from SARS-CoV-2 infected HBEC scRNA-seq, which was composed of 4,745 infected and 50,297 bystander cells (*39, 77*). The trained model reported an accuracy of 88%, a sensitivity of 62% and a specificity of 91%. In order to identify the importance of genes in the model’s prediction, we computed exact Shapley values for each prediction and genes using TreeSHAP (*78*). The genes were ranked by averaging the absolute Shapley values. Distribution of absolute Shapley values of genes identified by CRISPR screening were compared to the remaining other genes by a Wilcoxon rank sum test, and were found to be significantly higher ranked under the null hypothesis (p < 0.0001).

### Pseudovirus production

VSV-based pseudotype viruses were produced as described previously (*41*). Vector pCAGGS containing the SARS-Related Coronavirus 2, Wuhan-Hu-1 Spike Glycoprotein Gene, NR-52310, was produced under HHSN272201400008C and obtained through BEI Resources, NIAID, NIH. Briefly, 293T cells were transfected with pCAGGS vector expressing the SARS-CoV-2 spike glycoprotein and then inoculated with a replication-deficient VSV vector that contains expression cassettes for Renilla luciferase instead of the VSV-G open reading frame. After an incubation period of 1 hour at 37°C, the inoculum was removed and cells were washed with PBS before media supplemented with anti-VSV-G clone I4 was added in order to neutralize residual input virus (no antibody was added to cells expressing VSV-G). Pseudotyped particles were harvested 24 hour post inoculation, clarified from cellular debris by centrifugation and stored at −80°C before use.

### Pseudovirus entry assay

2 x 10^4^ Vero-E6 cells were seeded in 100 μl total volume each well of a black-walled clear bottom 96-well plate. The following day SARS-CoV-2 spike expressing VSV pseudovirus was added at 1:10 final concentration volume/volume and incubated for one day. Cells were lysed with Renilla-Glo or Renilla Luciferase Assay System (Promega) according to manufacturer instructions. The luciferase activity was measured using a microplate reader (Cytation 5, BioTek).

### Secondary assessment of CRISPR screen hits by cell viability

*HMGB1* sgRNAs were cloned into lentiCRISPRv2 (Addgene 52961) which also encodes the Cas9 gene (*79*). sgRNAs for all other genes were cloned into lentiGuide-Puro or a variant thereof, pXPR_050 (Addgene 52963, 96925, respectively). sgRNAs target sequences are in supplementary Table 1. Vero-E6-Cas9 cells were individually transduced with lentiviruses expressing one to three unique sgRNA per gene and then selected with puromycin for 7 days. After selection, 1.25 x 10^3^ cells were seeded in each well of a 384-well black walled clear bottom plate in 20 μl of DMEM + 5% FBS. The following day, 5 μl of SARS-CoV-2 was added for a final MOI of 0.1. Cells were incubated for three days before assessing cellular viability by CellTiter Glo (Promega). For each cell line, viability was determined in SARS-CoV-2 infected relative to mock infected cells. Five replicates per condition were performed in each of three independent experiments.

### SARS-CoV-2 fluorescent reporter virus assay

Cells were plated at 2.5 x 10^3^ cells per well in a 384-well plate and then the following day, icSARS-CoV-2-mNG was added at a MOI of 1.0. Infected cell frequencies as measured by mNeonGreen expression were assessed at 2 dpi by high content imaging (Cytation 5, BioTek) configured with bright field and GFP cubes. Total cell numbers were quantified by Gen5 software of brightfield images. Object analysis was used to determine the number of mNeon Green positive cells. The percentage of infection was calculated as the ratio between the number of GFP+ cells and the total amount of cells in brightfield. Data are normalized to the average of DMSO treated cells.

### Identification of anti-viral drugs targeting CRISPR gene hits

Calpain Inhibitor III (#14283), SIS3 (#15495), APY0201 (#9001589), and PFI-3 (#15267) were purchased from Cayman Chemical. Drugs were resuspended at a stock concentration of 40mM in DMSO and then two-fold serial dilutions were performed in DMSO. 20 nanoliters of 1000X drug stock were spotted into each well of a 384-well plate using Labcyte ECHO acoustic dispenser at the Yale Center for Molecular Discovery. 1.25 x 10^3^ Vero-E6 cells were plated per well in 20 μl of phenol-red free DMEM containing 5% FBS. Two days later, 5,000 PFU (MOI ~1) icSARS-CoV-2-mNG in 5 μl media was added. Cells were incubated at 37°C and 5% CO2 for two days. Infected cell frequencies were quantified by mNeonGreen at 2 dpi (Cytation 5, BioTek). In parallel, 500 PFU (MOI~0.2) SARS-CoV-2 in 5 μl media was added to replicate plates and cell viability was quantified by CellTiter Glo at 3 dpi.

### Western blot

Vero-E6 (1 x 10^6^) were collected and lysed in Nonidet P-40 lysis buffer (20 mM Tris-HCl [pH 7.4], 150 mM NaCl, 1 mM EDTA, 1% Nonidet P-40, 10 mg/ml aprotinin, 10 mg/ml leupeptin, and 1 mM PMSF). The cell lysates were fractionated on SDS-PAGE, and transferred to a PVDF membrane. Immunoblotting analyses were performed with anti-HMGB1 (Abcam 18256) and visualized with horseradish peroxidase-coupled goat anti-mouse/rabbit IgG using the Chemiluminescence Detection system.

---

**Fig. S1.**
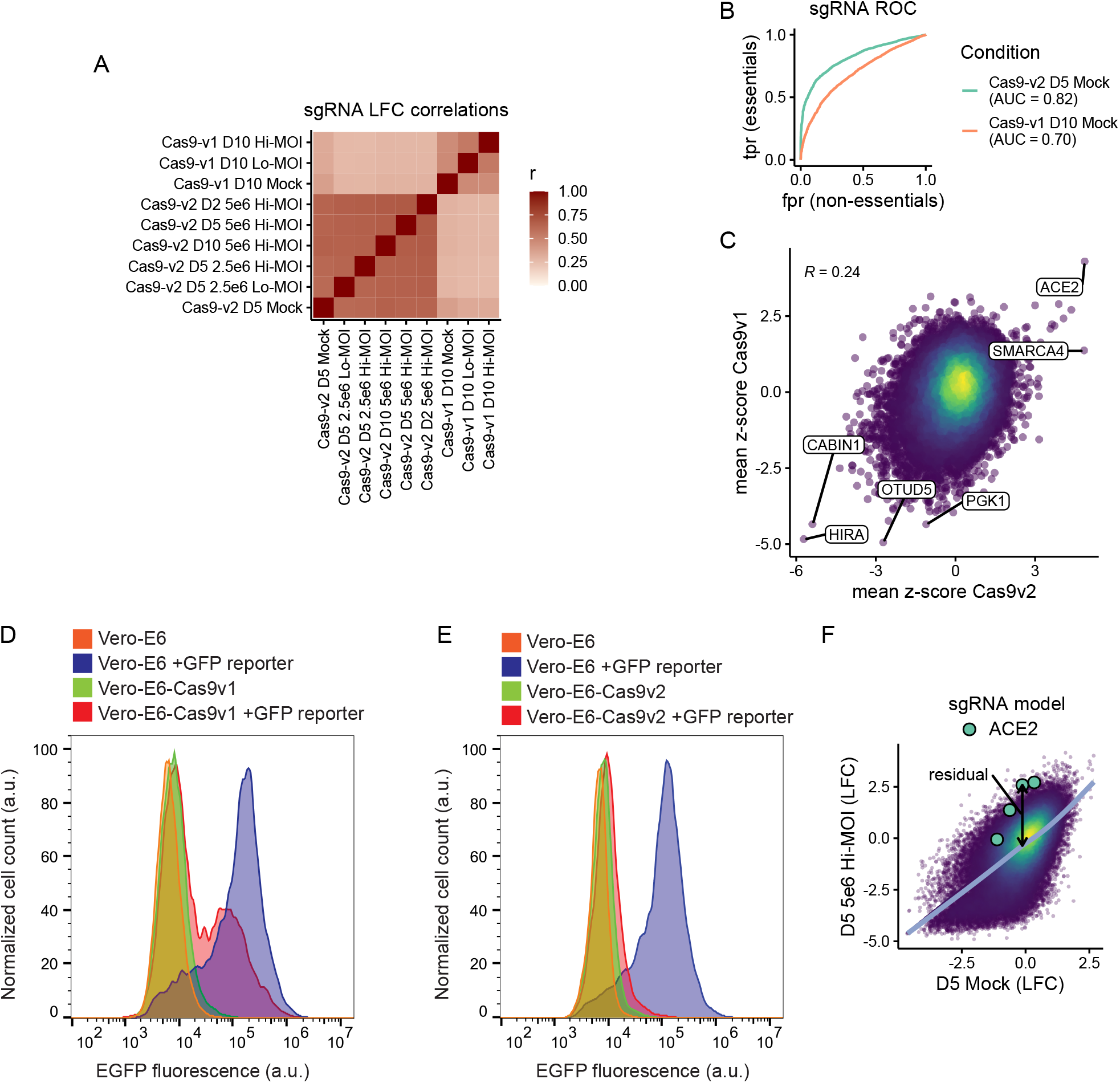
Quality control metrics for CRISPR screen. (A) Correlation matrix depicting the Pearson correlation between the guide-level log-fold change values relative to the plasmid DNA. Cells were cultured in DMEM with 2% FBS (D2), 5% FBS (D5), 10% FBS (D10), plated at 2.5 x 10^6^ or 5.0 x 10^6^ cells per flask and infected at a MOI 0.1 (Hi) or MOI 0.01 (Lo). (B) Receiver-operator characteristic (ROC) curve for the recovery of guides targeting essential genes in the mock-treated condition of the Cas9-v1 and Cas9-v2 screens. True positives are n = 1,528 essential genes (n = 6,178 guides); true negative genes are n = 622 non-essential genes (n = 2,504 guides). We mapped essential and non-essential genes, which were derived for human cell lines, to the African green monkey genome simply by matching gene symbols. AUC = area under curve. (C) Correlation between gene enrichment in Cas9-v1 and Cas9-v2 screens. Pearson correlation is reported. (D-E) GFP-based Cas9 activity assay in Vero-E6 cells stably expressing either Cas9-v1 (D) or Cas9-v2 (E). The pXPR_047 construct expresses GFP and an sgRNA targeting GFP; therefore, cells without Cas9 activity will express GFP, whereas cells with high Cas9 activity will knock out GFP and resemble parental cells. (F) Approach to calculate residuals from log-fold change data, using ACE2 and the 5%FBS, 5 x 10^6^ cells/flask, MOI 0.1 condition as an example. A natural cubic spline with four degrees of freedom is shown in blue, and a residual for each sgRNA is calculated to be the vertical distance from the fit spline.

**Fig. S2.**
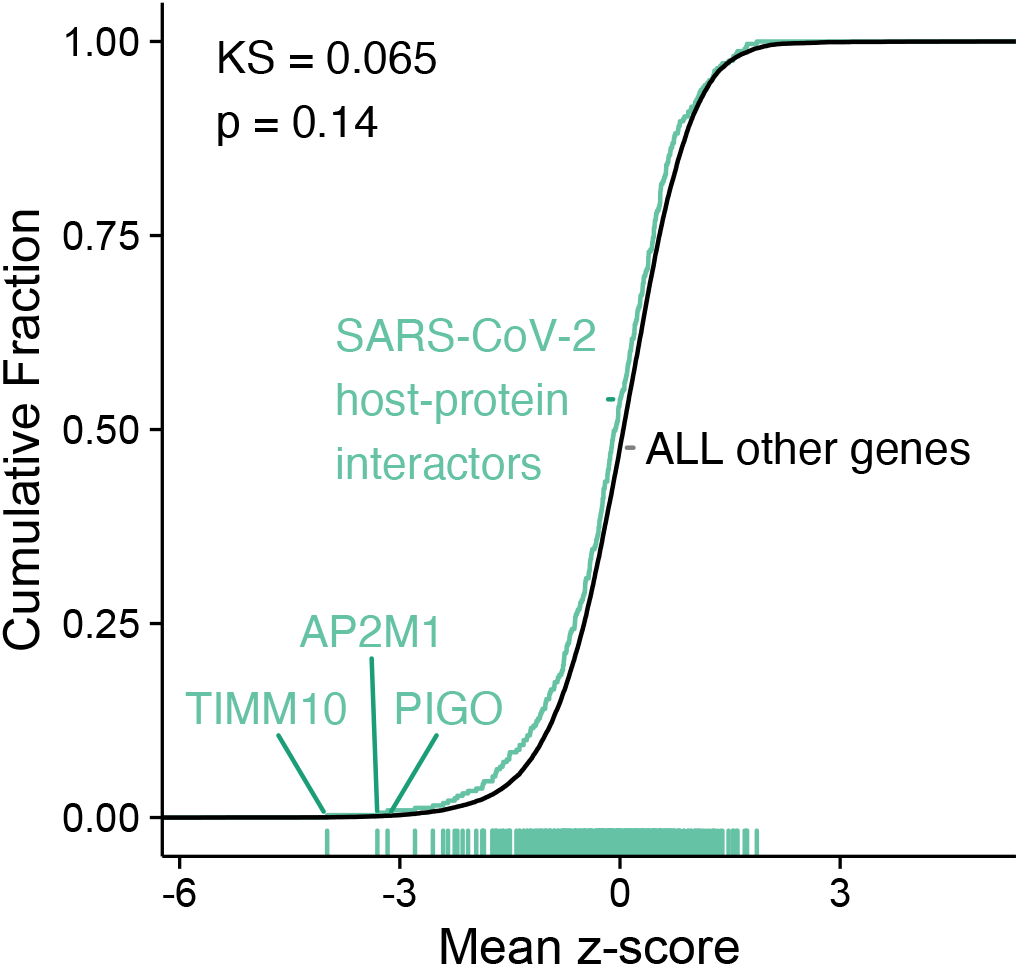
Comparison to SARS-CoV-2 protein interactome. Cumulative distribution of mean z-scores for SARS-CoV-2 protein interactors from (7) (n=321; green) compared with all other genes (n = 21,351; black). Interactors with an absolute z-score greater than 3 are labelled. The KS-statistic and corresponding p-value are indicated for a two-sided KS-test. We mapped interactors to the African Green Monkey genome by matching gene symbols.

**Fig. S3.**
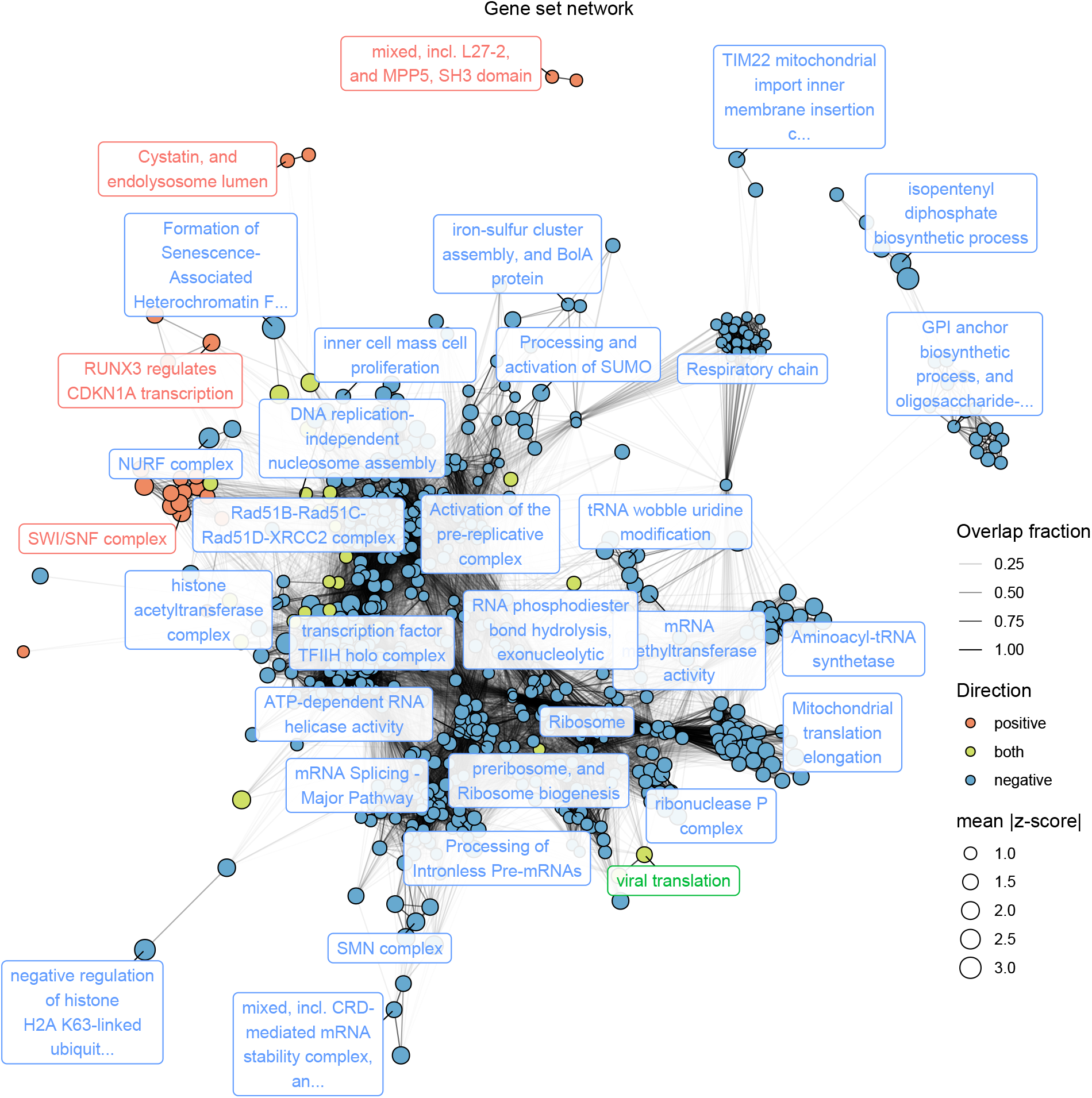
Network of gene sets. Nodes represent significantly enriched gene sets. The size of each gene set is proportional to its mean absolute z-score. Gene sets are colored by the direction in which they score. Edges represent significant overlap between gene sets. The transparency of each edge is proportional to the fraction of genes shared by two gene sets. Gene sets were clustered using the infomap algorithm and the most central set by PageRank is labelled for each cluster. The Fruchterman–Reingold algorithm was used to lay out the network.

**Fig. S4.**
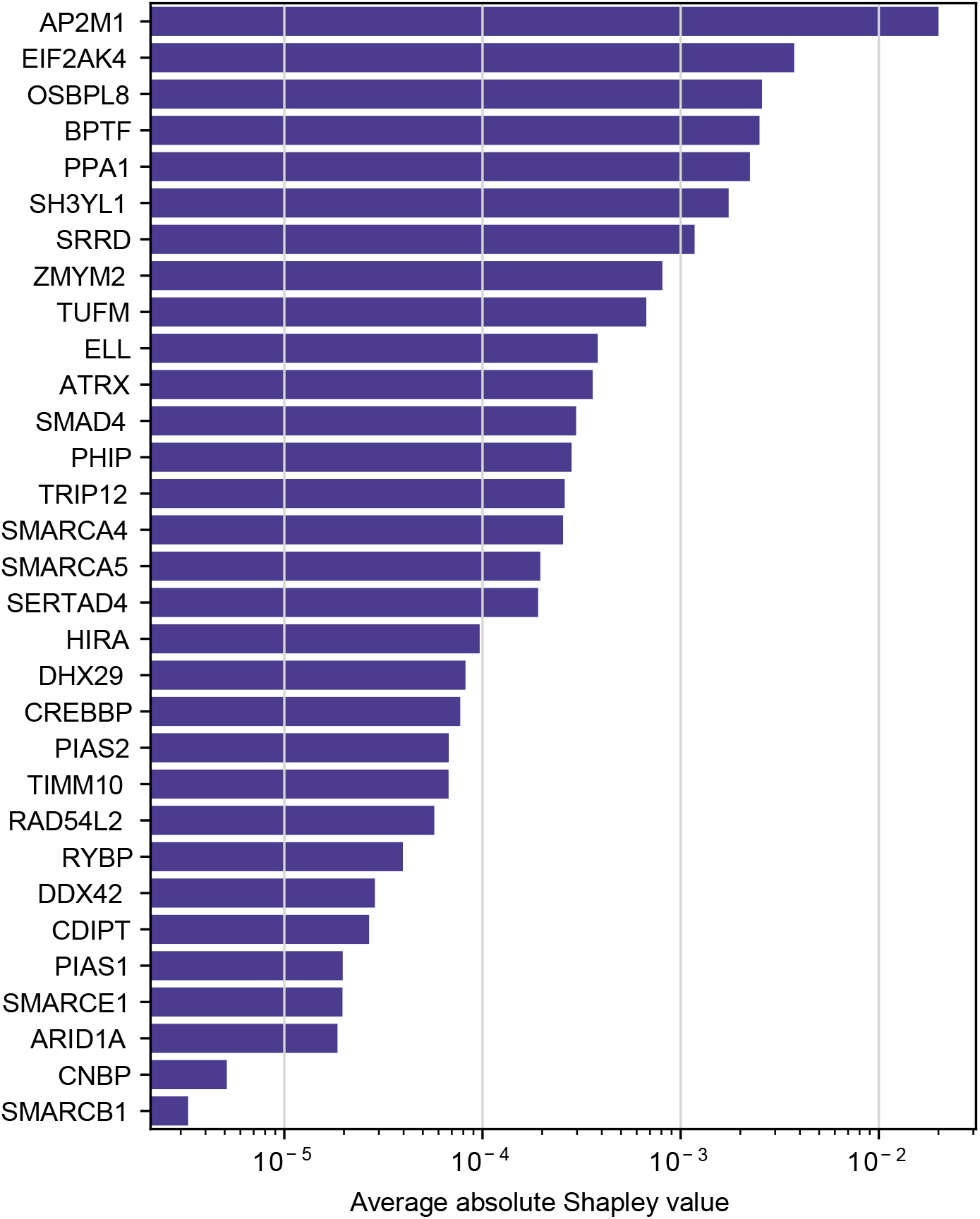
Feature importance of an infection prediction model trained on single-cell RNA sequencing analysis of infected human bronchial epithelial cells. Genes are ranked by mean of absolute Shapley values, computed from a gradient boosting model.

**Table S1.**
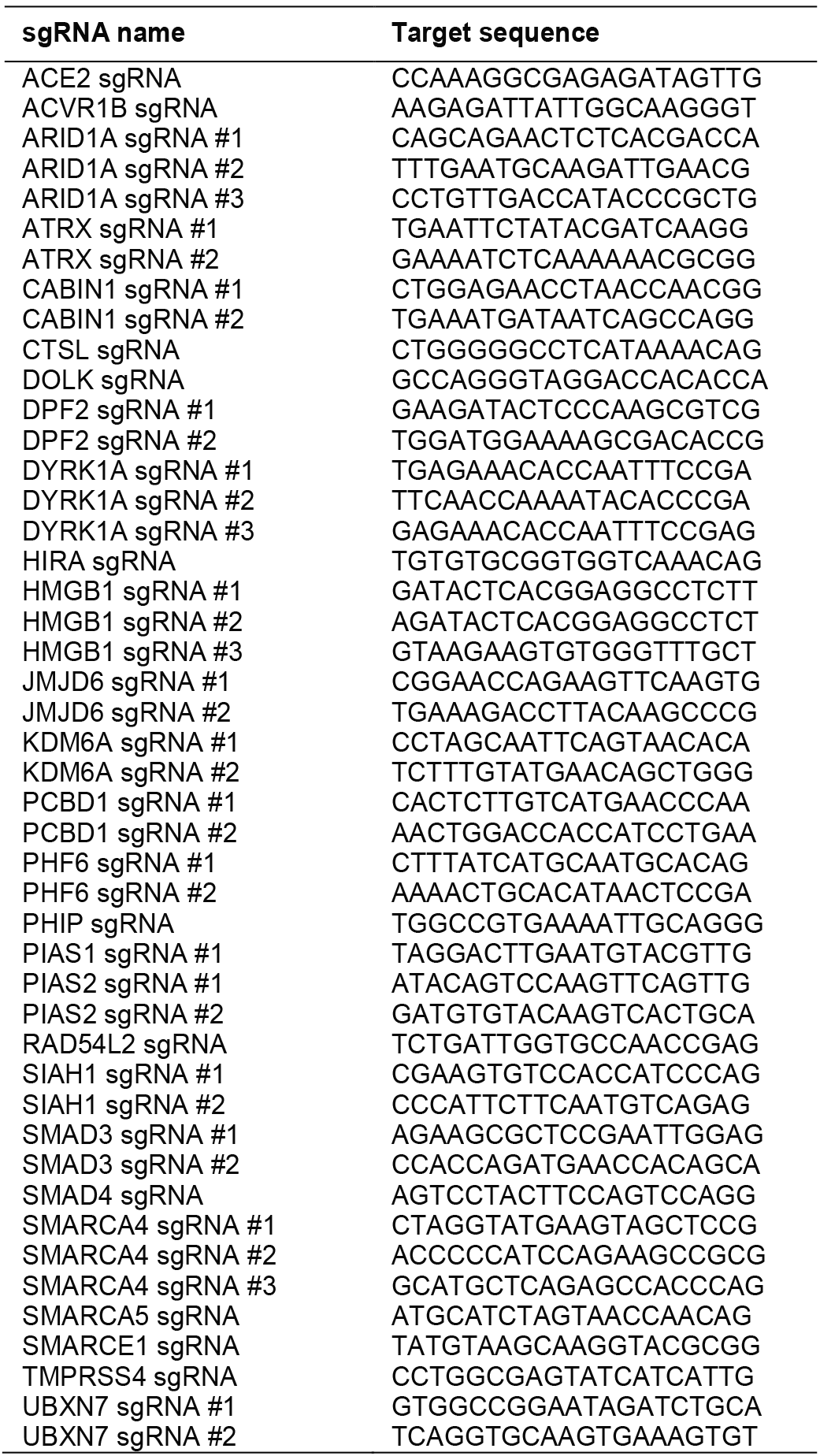
sgRNAs for secondary validation genes.

